# Parasites and invasions: changes in gastrointestinal helminth assemblages in invasive and native rodents in senegal

**DOI:** 10.1101/053082

**Authors:** Christophe Diagne, Alexis Ribas, Nathalie Charbonnel, Ambroise Dalecky, Caroline Tatard, Philippe Gauthier, Voitto Haukisalmi, Odile Fossati-Gaschignard, Khalilou Bâ, Mamadou Kane, Youssoupha Niang, Mamoudou Diallo, Aliou Sow, Sylvain Piry, Mbacké Sembène, Carine Brouat

**Author notes:** **Correspondence:** Christophe Diagne. E-mail address), Postal address: CBGP, 755 avenue du campus Agropolis, 34988 Montferrier-sur-Lez. /.

## Abstract

Understanding why some exotic species become widespread and abundant in their colonized range is a fundamental issue that still needs to be addressed. Among many hypotheses, newly established host populations may benefit from a parasite loss (“enemy release” hypothesis) through impoverishment of their original parasite communities or reduced infection levels. Moreover, the fitness of competing native hosts may be affected by the acquisition of exotic taxa from invaders (“parasite spillover”) and/or by an increased transmission risk of native parasites due to their amplification by invaders (“parasite spillback”). We focused on gastrointestinal helminth communities to determine whether these predictions could explain the ongoing invasion success of the commensal house mouse (*Mus musculus domesticus*) and black rat (*Rattus rattus*), as well as the associated drop of native Mastomys species, in Senegal. For both invasive species, our results were consistent with the predictions of the enemy release hypothesis. A decrease of helminth overall prevalence and individual species richness was observed along the invasion gradients as well as lower specific prevalence/abundance (*Aspiculuris tetraptera in M. m. domesticus, Hymenolepis diminuta in R. rattus*) on the invasion fronts. Conversely, we did not find strong evidence of helminth spill-over or spill-back in invasion fronts, where native and invasive rodents co-occurred. Further experimental research is needed to determine whether and how the loss of helminths and reduced infection levels along invasion routes may result in any advantageous effects on invader fitness and competitive advantage.

## 1. Introduction

Parasites *sensu lato* are likely to have a strong influence on their host population ecology, evolution and dynamics by exerting strong selection pressures on host life-history traits (Phillips *et al.* 2010). Over the last decade, parasitism has been considered as a key factor underlying expansion success of some invading organisms, especially in animal invasions (Prenter *et al.* 2004). Three major and not mutually exclusive mechanisms have been emphasized. The most evocative is the enemy release hypothesis, which states that invasive species may benefit in their introduced range from the escape of their natural parasites (Keane & Crawley 2002; Torchin et al. 2003; Mitchell & Power 2003; Colautti *et al.* 2004). The loss or reduced infection levels of parasite(s) may enhance host fitness and performances in the new environment compared to the original range, then facilitating its settlement and spread. Support for the enemy release hypothesis in the course of invasion has come from both metaanalysis (e;g., Torchin et al. 2003) and individual empirical studies previously conducted in diverse taxa (plants, invertebrates, and vertebrates) (e.g., Menendez *et al.* 2008, Phillips et al. 2010, Flory *et al.* 2011). The spillover hypothesis states that exotic hosts may introduce some of their coevolved parasites, these latter having negative impacts on native hosts of the introduction area (Tompkins et al. 2002; Prenter et al. 2004; Bell et al. 2009). Finally, the spillback hypothesis states that invaders may be competent hosts for local parasites, leading to increased density and transmission of infective stages in environment at the expense of native hosts (Kelly *et al.* 2009). Despite a burgeoning interest on parasitism-related invasion processes in the scientific literature, several gaps were recently highlighted (Heger & Jeschke 2014), especially on animal models. First, only some studies on invertebrates relied on the prior identification of invasion pathways (Wattier et al. 2007; Slothouber Galbreath et al. 2010; Lester et al. 2014), although this step has been recognized as critical to design reliable sampling strategies and robust comparative analyses in invasion research (Muirhead *et al.* 2008). Also, several studies focused only on invasive host species and their parasite communities, assuming often that native parasite communities are unimportant in invasion success (Kelly *et al.* 2009). Studies on invertebrates (Mastitsky et al. 2010; Slothouber Galbreath et al. 2010; Rode et al. 2013; Jones & Brown 2014) or amphibians (Shine, 2014) have however shown that information on native communities is crucial to distinguish spillover or spillback processes.

We propose to investigate parasitism-invasion success relationships by considering native and invasive host and parasite communities along two invasion gradients. We focused on the ongoing invasions of two major exotic species, the black rat (*Rattus rattus*) and the house mouse (*Mus musculus domesticus*) in Senegal (West Africa). Originating from Asia, *R. rattus* and *M. m. domesticus* made use of human migrations to expand their distribution range worldwide (Aplin *et al.* 2011; Bonhomme *et al.* 2011). Several studies have documented dramatic parasite-mediated impacts of these invasive rodents on insular indigenous faunas (e.g., Wyatt *et al.* 2008; Harris 2009), suggesting that they may be suitable biological models to study the relationships between parasitism and invasion success In Senegal, both taxa are exclusively commensal, with distribution areas covering now much of North and Central Senegal for *M. m. domesticus* and much of Senegal South of Gambia River for *R. rattus* (Figure 1). Historical records (see references in Konecny *et al.* 2013; Dalecky *et al.* 2015) and molecular analyses (Konecny *et al.* 2013; Lippens *et al.* in revision) showed that these rodents were first brought to Senegalese coasts by European explorers and settlers, and remained in coastal villages and towns until the beginning of the 20^th^ century. Both taxa have spread further inland during the last century via well-defined invasion routes, thanks to the improvement of human activities and transport infrastructures, and resulting in the extirpation of native rodents (mostly *Mastomys* species) from commensal habitats beyond their invasion front (Dalecky *et al.* 2015).

**Figure 1.**
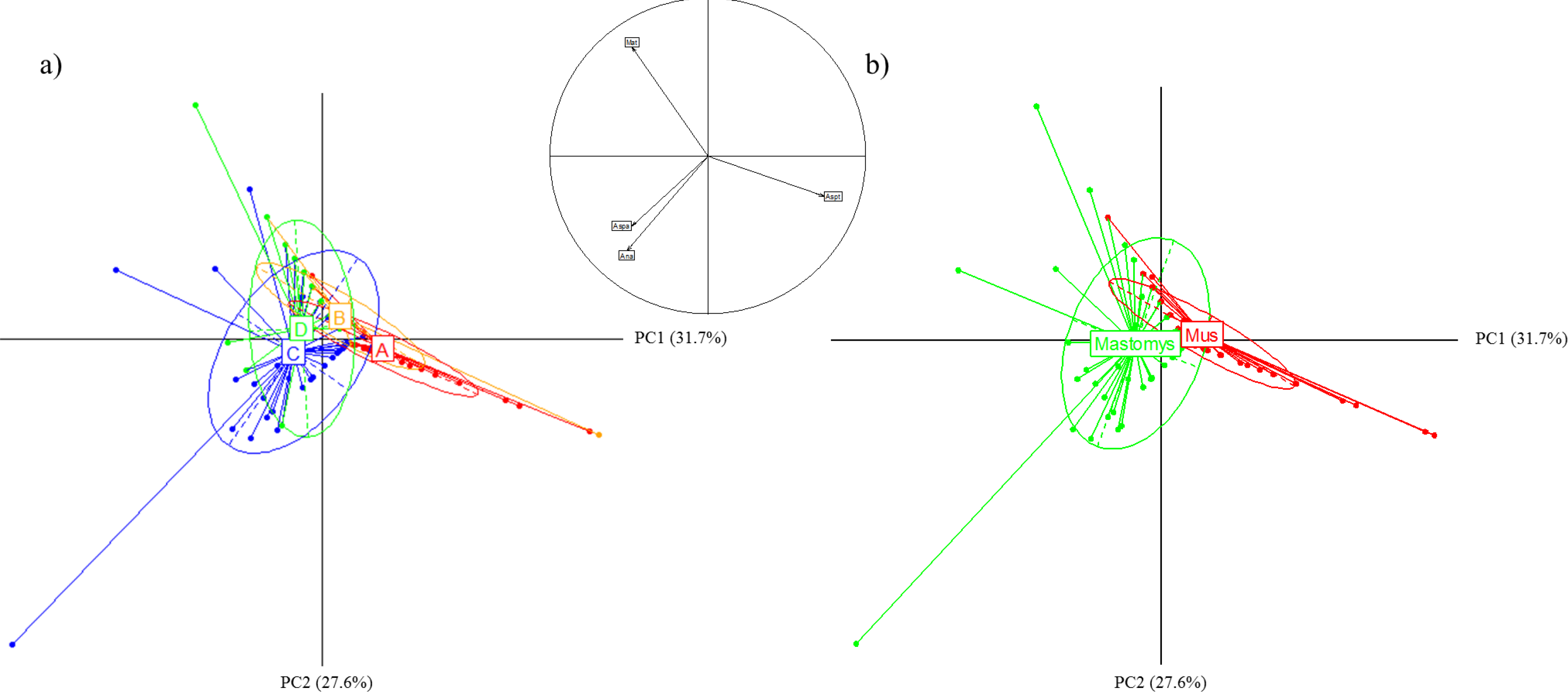
Rodent sampling sites on house mouse (symbols in white) and black rat (symbols in black) invasion routes. Locality codes are given in Table 1. Triangles, squares and circles correspond respectively to sites of long-established invasion, recently invaded sites and noninvaded sites.

We focused on gastrointestinal helminths communities in commensal rodents from Senegal to test some predictions relating animal invasion success and parasitism. Besides of their known regulatory effects on rodent fitness (Deter *et al.* 2007) and population dynamics (Vandegrift & Hudson 2009), gastro-intestinal helminths are highly diversified and prevalent in rodent populations in Senegal (Brouat *et al.* 2007). Using an integrative framework combining systematics and community ecology, we compared gastro-intestinal helminth communities in natural populations of native and exotic rodents along well-described invasion routes in Senegal. Under the enemy release hypothesis, we expected to detect a decrease of parasitism in exotic rodents along their invasion route. The focus on Senegalese populations of exotic rodents was assumed, as parasitological signatures from their putative European sources have most probably disappeared during the last centuries. On the other hand, we looked for evidence of parasites common to native and exotic rodents in newly invaded areas. Under the spillover and spillback hypotheses, we expected to detect an increase of parasitism related to these common helminths in native rodents from invaded compared to those from non-invaded areas. The consideration of non-invaded areas in the comparative design allowed having an overview of helminth assemblages infecting native rodents before the arrival of exotic hosts.

## 2. Materials and Methods

### 2.1 Ethical statement

Trapping campaigns within private properties was systematically realized with prior explicit agreement from relevant institutional (CBGP: 34 169 003) and local authorities. All animal-related procedures were carried out following official ethical guidelines (Sikes, Gannon & Amer Soc 2011).

### 2.2 Rodent sampling

In Senegal, the distributions of exotic rodents are mainly restricted to villages and towns (Granjon & Duplantier2009). Recent molecular analyses have enabled to trace the invasion routes of *M. m. domesticus* (Lippens *et al.* in revision) and *R. rattus* (Konecny *et al.* 2013) within the country. On the basis of these studies, sampling sites have been chosen along one invasion route for each exotic rodent: in the Senegal River valley for *M. m. domesticus,* along which populations firstly introduced in the region of St Louis have spread East (Lippens *et al.* in revision); in the southern part of the country for *R. rattus,* where eastern populations are from the genetic group at the origin of recently introduced populations in southwestern Senegal (Konecny *et al.* 2013) (Fig. 1). Three categories of sites describing the invasion status were sampled along each invasion route: (i) sites of long-established invasion on the west coast, where invasive rodents have settled since centuries and are highly dominant or even the single commensal species present; (ii) recently invaded sites (i.e. invasion front), where invasive rodents have settled for less than 30 years and occur in sympatry with native rodent species; and (iii) non-invaded sites, where invasive rodents have never been recorded. For each category of invasion status, three to six sites were sampled (Fig. 1).

Fieldwork was conducted inside human dwellings in March-April 2013 for *M. m. domesticus* and from November 2013 to February 2014 for *R. rattus.* The detailed description of the standardized rodent trapping protocol used here was provided in Dalecky *et al.* (2015). Briefly, we set at least 120 traps (two traps per house, with sampled houses chosen to cover a significant part of the inhabited area) during one to three nights, in order to ensure that 20 adult individuals per rodent species were caught in each locality. Rodents were captured alive and sacrificed by cervical dislocation, weighted to the nearest 0.5 g, sexed and dissected. Finally, digestive tracts were removed, unrolled and individually stored until examination in plastic universal vials containing 95% ethanol. Rodents were aged on the basis of body weight and/or reproductive status following Granjon and Duplantier (2009). They were identified with morphometric and genetic tools (cytochrome b gene-based RFLP for specific identification in *Mastomys* spp.; ZFY2 gene-based RFLP for subspecific identification in M *musculus*).

### 2.3 Collection and identification of gastrointestinal helminths

For each rodent, the different sections of the digestive tract (stomach, small intestine, large intestine and caecum) were scrutinized following Ribas *et al.* (2011). Helminths were carefully removed and counted, then classified by morphotype and stored in 95% ethanol. For accurate identification at the most precise taxonomic level, we combined morphological and molecular approaches as diagnosis tools. Morphological identification was firstly carried out using conventional microscopy and generalist identification keys (Khalil, Jones & Bray 1994; Anderson, Chabaud & Willmott 2009) or specific literature when available. At least one specimen of each taxon identified per rodent species and locality category was then sequenced for Cytochrome Oxidase 1 (CO1) for nematodes (Cross *et al.* 2007) and acanthocephalan, and Nicotinamide Adenine Dinucleotide subunit 1 (NAD1) for cestodes (Littlewood, Waeschenbach & Nikolov 2008). For this purpose, total DNA was extracted from the midbody region of the individual helminth, with the anterior and posterior regions retained in 95% ethanol to complete the morphological examination if necessary. DNA extraction was achieved using the DNeasy blood and tissue Kit (Qiagen) according to manufacturer’s instructions slightly modified with a final elution of 200μl of AE buffer. Tissue samples were digested in 180μl of lysis buffer with 20μl of proteinase-K incubated at 56°C overnight. PCR amplifications were performed using the primers 5’-TTGRTTTTTTGGTCATCCTGARG-3’ and 5’ -W S YM AC W AC AT A AT AAGTAT C ATG-3’ for CO1, and 5’-GGNTATTSTCARTNTCGTAAGGG-3’ and 5’-TTCYTGAAGTTAACAGCATCA-3’ for NAD1, in 25 μl reactions containing 2 μl of DNA extract, 1X of Dream Taq buffer (included 2mM of MgCl2), 0.2 mM of dNTP, 0.5 μM of each primer and 1 Unit of Dream Taq (ThermoFischer). Cycling conditions in Mastercycler gradient (Eppendorf) were the following: 94 °C 3 min, followed by 40 cycles of 94 °C 30s, 50 °C 60s for CO1 and 55°C 60s for NAD1, 72 °C 90s, and a final extension at 72 °C 10 min. PCR products were run on a 1.5% agarose gel to ensure amplification and then sequenced in both direction by Eurofins MWG (Germany). Sequences obtained were processed (cleaning, assembling and alignment) then compared to both public (Genbank) and personal molecular databases. The personal reference sequence database was developed by sequencing nematodes, cestodes and acanthocephalans of rodents from different West-African areas (Burkina Faso, Mali, Senegal; Supplementary Fig. S1) that had been already identified morphologically to the species level.

### 2.4 Data analysis

The analyses were carried out independently for each invasion route.

*Structure of helminth assemblages.* Using the software Quantitative Parasitology 3.0 (Rozsa, Reiczigel & Majorost 2000), prevalence (i.e., percentage of infected hosts) and mean abundance of each helminth were estimated per host species in each locality (Table 2). We also investigated whether the helminth community was structured according to host species and/or invasion status along each invasion route. We thus performed a Principal Component Analysis (PCA) on the restricted dataset including infected hosts only and the presence/absence of GIH taxa showing prevalence higher than 5%. The significance of host species and invasion status was tested independently using Between/Within-groups Analysis (BWA) and Monte-Carlo tests (999 permutations). Note that the influence of invasion status was analyzed by considering four groups to avoid a host-species bias in recently invaded sites: A) invasive hosts in sites of long-established invasion, B) invasive hosts in sites at the invasion front, C) native hosts in sites at the invasion front and D) native hosts in non-invaded sites.

*Testing factors affecting helminth assemblages.* We used Generalized Linear Models (GLMs) to evaluate whether variations in helminth communities along each invasion route were consistent with the hypotheses relating parasitism and invasion success. We conducted separate analyses for native and invasive host species as we expected different outcomes for these rodents:

1. In exotic rodents, the examination of the enemy release hypothesis was performed using the following response variables calculated for each individual host: overall prevalence (presence/absence of helminths, combining all taxa), species richness (number of helminth taxa in one individual host), specific prevalence (presence/absence of a given helminth taxon) and specific abundance (number of individuals of a given helminth taxon). These variables were expected to decrease from sites of long-established invasion to invasion fronts under the enemy release hypothesis (e.g., Wattier *et al.* 2007).
2. In native rodents, the examination of spillover and spillback hypotheses was performed using species richness and specific abundance and prevalence of helminths common to native and exotic rodents as response variables. We expected to detect an increase of species richness at invasion fronts compared to not invaded sites under both hypotheses, and an increase of specific prevalence or abundance of native or exotic helminths under the spillback or spillover hypotheses, respectively. We had no particular expectation on overall helminth prevalence and abundance.

For specific indicators, only helminth taxa that exhibited prevalence levels higher than 10% were considered. We assumed a binomial distribution for prevalence data and a Poisson distribution for abundance data and species richness, using then respectively quasibinomial and negative binomial distributions in case of overdispersion. The full models included individual host factors (sex and body mass), the invasion status of the locality (long-established invasion *vs* invasion front for invasive hosts, invasion front *vs* not invaded sites for native hosts) and their pairwise interactions as possible predictors. As some helminths of terrestrial mammals spend at least one part of their life-cycle in the external environment as egg or larvae, we included the climate as environmental predictor at the scale of the locality. For this purpose, we first carried out PCA on climatic data covering the period between 1997 and 2012 (temperature recorded from local weather stations closest to sampled localities and available on http://www.ncdc.noaa.gov/cdo-web/datasets;; rainfall: recorded from satellite products available on http://richardis.univ-paris1.fr/precip/rainser1.html with GPCP-1DD as date source; ; Supplementary Fig. S2, Fig. S3). Then, we included the first PCA axis coordinates of each locality in the starting model. If strong association was graphically observed between the first PCA axis and invasion status, the coordinates on the second axis were included. Model selection was then performed from full models based on the Akaike information criterion with correction for samples of finite size (AICc). The most parsimonious model among those selected within two AIC units of the best model was chosen. *P-values* were obtained by stepwise model simplification using likelihood-ratio tests. The final models were validated by checking the model dispersion and ensuring normality, independence and variance homogeneity of the residuals. All analyses were performed using the packages ade4 v1.4-16, MuMIn v1.15.1 and lme4 v1.1-8 implemented in R software v3.2.1 (R Core Team 2015).

## 3. Results

### 3.1 Rodent rapping

A total of 752 rodents belonging to four species (268 *M. m. domesticus* and 169 *Mastomys erythroleucus* on the mouse invasion route; 193 *R. rattus,* 29 *Ma. erythroleucus* and 93 *Mastomys natalensis* on the rat invasion route) were collected in 25 sites (Table 1). In sites of long-established invasion, only invasive rodents were generally captured except on the rat invasion route where few *Ma. erythroleucus* individuals (n = 24) have been trapped but not included in further analyses due to low sample size. As expected at the invasion front, native rodents co-occurred with invasive ones, although being nearly systematically less abundant. Along the mouse invasion route, native rodent communities were largely dominated by *Ma. erythroleucus,* whereas *Ma. erythroleucus* and *Ma natalensis* dominated along the rat invasion route. These two native sister species did not co-occur, except in a single non-invaded locality (Bransan). We therefore did not disentangle the potential impact of these *Mastomys* sibling species in the analyses focusing on the rat invasion route.

**Table 1:**
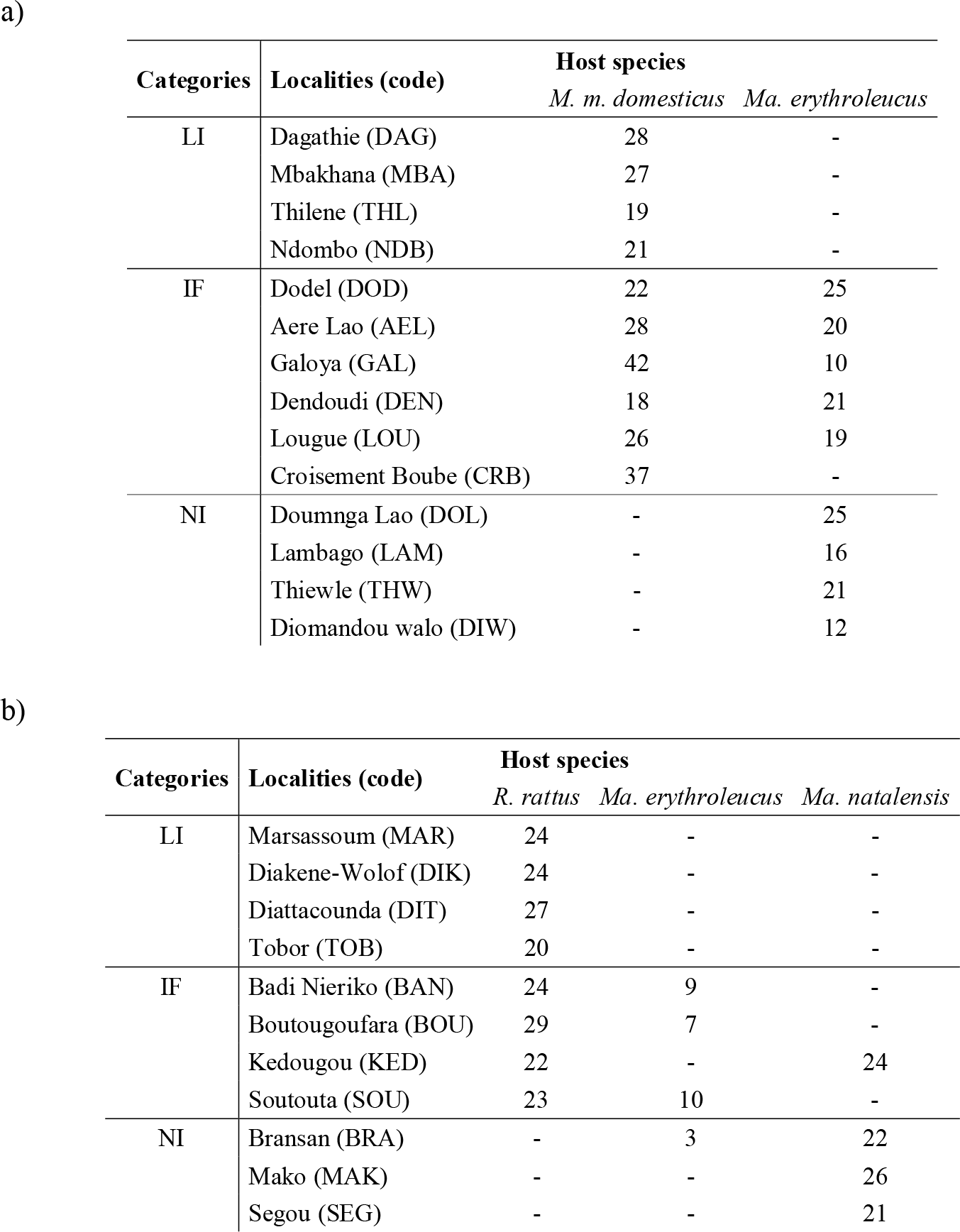
Number of individuals analyzed per host species for each locality along a) the mouse invasion route and b) the rat invasion route. The code used for each sampling locality is indicated in parentheses. LI: localities of long-established invasion; IF: localities at invasion fronts; NI: non-invaded localities. ‘-’ indicates that no rodent was trapped or analyzed.

### 3.2 Structure of helminth assemblages

We recorded eight taxa of helminth along the mouse invasion route (Table 2). Five nematode taxa (*Aspiculuris tetraptera, Gongylonema* sp., *Pterygodermatites senegalensis, Syphacia obvelata*, Trichostrongylid) were found in *M. m. domesticus* only, and two species (*Anatrichosoma* sp., *Asp. africana*) were found in *Ma. erythroleucus* only. The dominant *Asp. tetraptera* was the only nematode found in M *m. domesticus* populations from long-established and recently established invasion. *Aspiculuris africana* was restricted on native rodents from invasion front sites. Only the cestode *Mathevotaenia symmetrica* was found in both host species whatever the invasion status of the sampled locality. The overall prevalence varied from 0.3% (Trichostrongylid) to 11.6% (*Asp. tetraptera*) in *M. m domesticus,* and from 15.4% (*Asp. africana*) to 36.1% (*M. symmetrica*) in *Ma. erythroleucus.*

**Table 2:**
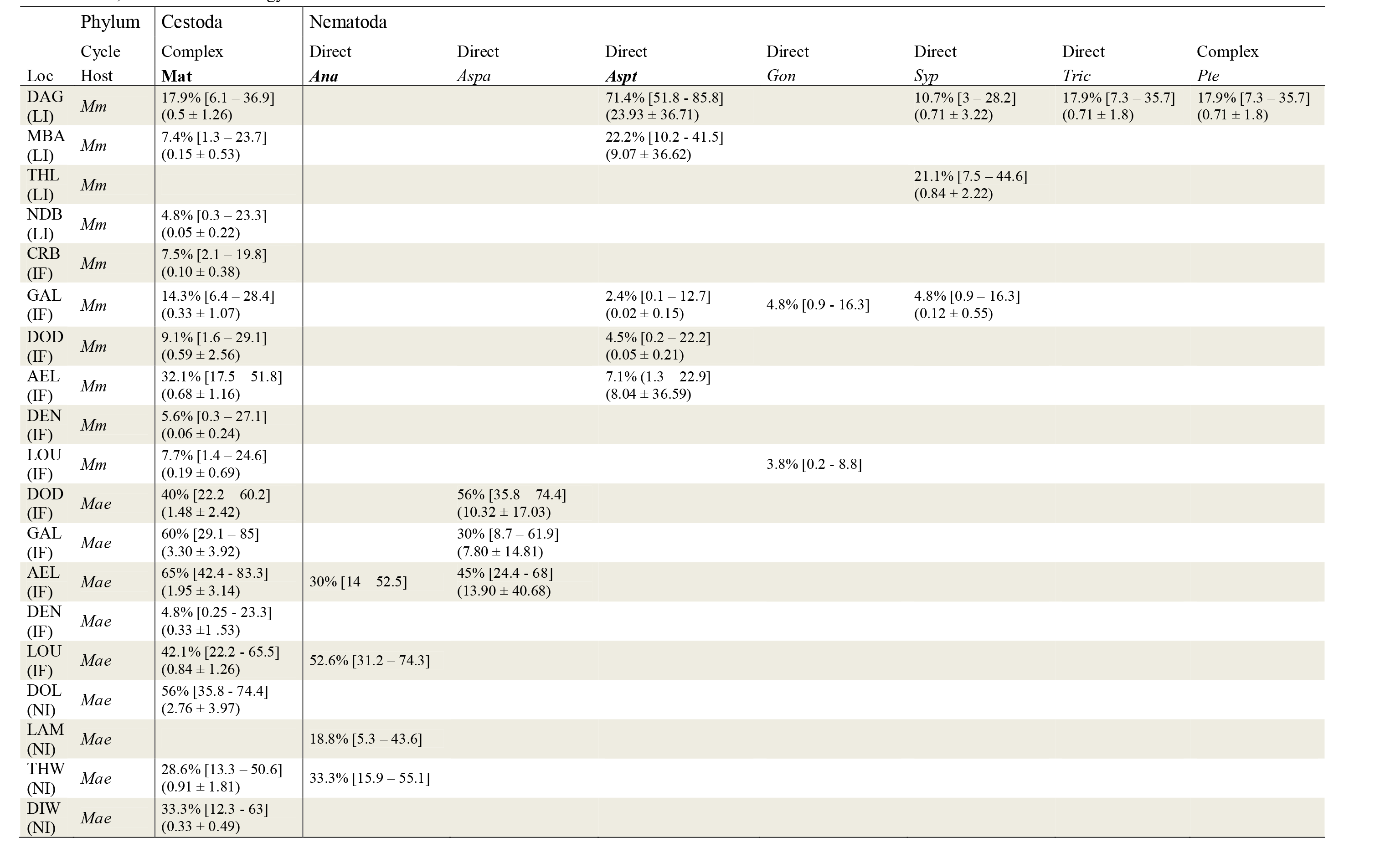
Prevalence in % [with 95% confidence intervals calculated with Sterne’s exact method] and abundances (mean ± standard deviation) of GIH taxa collected from *M. m. domesticus* (*Mm*) and *Ma. erythroleucus* (*Mae*) for each sampling locality (codes are provided in Table 1; LI: localities of long-established invasion; IF: invasion front; NI: non-invaded localities.) along the mouse invasion route. Taxa in bold are those chosen for performing GLMs. No abundance data was reported for both *Anatrichosoma* sp. and *Gongylonema* sp as they were difficult to quantify. The type of lifecycle, direct (only one host in the cycle) or complex (at least one intermediate host, mainly insects), is also provided for each GIH taxon. Legend: Loc = Locality. Cestoda: Mat = *Mathevotaenia symmetrica*. Nematoda: Ana = *Anatrichosoma* sp.; Aspa = *Aspiculuris africana*; Aspt = *Aspiculuris tetraptera*; Gong = *Gongylonema* sp.; Pte = *Pterygodermatites senegalensis*; Syp = *Syphacia* obvelata; Tric = Trichostrongylid.

| Loc | Phylum<br>Cycle<br>Host | Cestoda | Nematoda |  |  |  |  |  |  |
| --- | --- | --- | --- | --- | --- | --- | --- | --- | --- |
|  |  | Complex<br>Mat | Direct<br>Ana | Direct<br>Aspa | Direct<br>Aspt | Direct<br>Gon | Direct<br>Syp | Direct<br>Tric | Complex<br>Pte |
| DAG (LI) | *Mm* | 17.9% [6.1 – 36.9]<br>(0.5 ± 1.26) | | | 71.4% [51.8 - 85.8]<br>(23.93 ± 36.71) | | 10.7% [3 – 28.2]<br>(0.71 ± 3.22) | 17.9% [7.3 – 35.7]<br>(0.71 ± 1.8) | 17.9% [7.3 – 35.7]<br>(0.71 ± 1.8) |
| MBA (LI) | *Mm* | 7.4% [1.3 – 23.7]<br>(0.15 ± 0.53) | | | 22.2% [10.2 - 41.5]<br>(9.07 ± 36.62) | | | | |
| THL (LI) | *Mm* | | | | | | 21.1% [7.5 – 44.6]<br>(0.84 ± 2.22) | | |
| NDB (LI) | *Mm* | 4.8% [0.3 – 23.3]<br>(0.05 ± 0.22) | | | | | | | |
| CRB (IF) | *Mm* | 7.5% [2.1 – 19.8]<br>(0.10 ± 0.38) | | | | | | | |
| GAL (IF) | *Mm* | 14.3% [6.4 – 28.4]<br>(0.33 ± 1.07) | | | 2.4% [0.1 – 12.7]<br>(0.02 ± 0.15) | 4.8% [0.9 - 16.3] | 4.8% [0.9 – 16.3]<br>(0.12 ± 0.55) | | |
| DOD (IF) | *Mm* | 9.1% [1.6 – 29.1]<br>(0.59 ± 2.56) | | | 4.5% [0.2 – 22.2]<br>(0.05 ± 0.21) | | | | |
| AEL (IF) | *Mm* | 32.1% [17.5 – 51.8]<br>(0.68 ± 1.16) | | | 7.1% (1.3 – 22.9)<br>(8.04 ± 36.59) | | | | |
| DEN (IF) | *Mm* | 5.6% [0.3 – 27.1]<br>(0.06 ± 0.24) | | | | | | | |
| LOU (IF) | *Mm* | 7.7% [1.4 – 24.6]<br>(0.19 ± 0.69) | | | | 3.8% [0.2 - 8.8] | | | |
| DOD (IF) | *Mae* | 40% [22.2 – 60.2]<br>(1.48 ± 2.42) | | 56% [35.8 – 74.4]<br>(10.32 ± 17.03) | | | | | |
| GAL (IF) | *Mae* | 60% [29.1 – 85]<br>(3.30 ± 3.92) | | 30% [8.7 – 61.9]<br>(7.80 ± 14.81) | | | | | |
| AEL (IF) | *Mae* | 65% [42.4 - 83.3]<br>(1.95 ± 3.14) | 30% [14 – 52.5] | 45% [24.4 - 68]<br>(13.90 ± 40.68) | | | | | |
| DEN (IF) | *Mae* | 4.8% [0.25 - 23.3]<br>(0.33 ± 1.53) | | | | | | | |
| LOU (IF) | *Mae* | 42.1% [22.2 - 65.5]<br>(0.84 ± 1.26) | 52.6% [31.2 – 74.3] | | | | | | |
| DOL (NI) | *Mae* | 56% [35.8 - 74.4]<br>(2.76 ± 3.97) | | | | | | | |
| LAM (NI) | *Mae* |  | 18.8% [5.3 – 43.6] |  |  |  |  |  |  |
| THW (NI) | *Mae* | 28.6% [13.3 – 50.6]<br>(0.91 ± 1.81) | 33.3% [15.9 – 55.1] | | | | | | |
| DIW (NI) | *Mae* | 33.3% [12.3 - 63]<br>(0.33 ± 0.49) | | | | | | | |

We recorded 14 taxa of helminths in rodents sampled along the rat invasion route (Table 3). Three nematodes (*Aspiculuris* sp., *Gongylonema* sp., *Protospirura muricola*) and the acantocephalan *Moniliformis moniliformis* were strictly found within R. *rattus* sampled in sites of long-established invasion. Three nematodes (Asp. *africana, Pterygodermatites* sp., *Trichuris mastomysi*) and one cestode (*Raillietina baeri*) were specifically found in *Mastomys* spp. Two nematodes (*Neoheligmonella granjoni, Physaloptera* sp.) and four cestodes (*Hymenolepis diminuta, Hymenolepis sp., M. symmetrica, Raillietina trapezoides*) were found both in *R. rattus* and *Mastomys* spp., but only two species (N *granjoni., R. trapezoides*) were shared by sympatric invasive and native rodent populations at the invasion front. *Aspiculuris* sp. in *R rattus* would be a new species (Ribas et al.unpublished work). The overall prevalence ranged from 1% (*R. trapezoides*) to 30.1% (*H diminuta*) in *R. rattus,* and from 0.8% (*Pterygodermatites* sp., *H. diminuta*) to 26.2% (*T. mastomysi*) in *Mastomys* spp.

**Table 3:** Prevalence in % [with 95% confidence intervals calculated with Sterne’s exact method] and abundances (mean ± standard deviation) of GIH taxa collected from *R. rattus* (*Rr*), *Ma. erythroleucus* (*Mae*) and *Ma. natalensis* (*Man*) for each sampling locality (codes are provided in table 1; LI: localities of long-established invasion; IF: invasion front; NI: non-invaded localities.) along the rat invasion route. Taxa in bold are those chosen for performing GLMs. No abundance data was reported for *Gongylonema* sp. as it was difficult to quantify. The type of lifecycle, direct (only one definitive host) or complex (at least one intermediate host, mainly insects), is also provided for each GIH taxon. Legend: Loc = locality code. Acanthocephalan: Mon = *Moniliformis moniliformis*. Cestoda: Hym = *Hymenolepis* sp.; Hymd = *Hymenolepis diminuta*; Mat = *Mathevotaenia symmetrica*; Raib = *Raillietina baeri*; Rait = *Raillietina trapezoides*. Nematoda: Asp = *Aspiculuris* sp.; Aspa = *Aspiculuris africana*; Gong = *Gongylonema* sp.; Neo = *Neoheligmonella granjoni*; Phy = *Physaloptera* sp.; Pro = *Protospirura muricola*; Pte = *Pterygodermatites* sp.; Tri = *Trichuris mastomysi*.

| Loc | Phylum | Acanthocephala | Cestoda |  |  |  |  | Nematoda |  |  |  |  |  |  |  |
| --- | --- | --- | --- | --- | --- | --- | --- | --- | --- | --- | --- | --- | --- | --- | --- |
|  | Cycle | Complex | Complex | Complex | Complex | Complex | Complex | Direct | Direct | Direct | Direct | Complex | Complex | Complex | Direct |
|  | Host | *Mon* | *Hym* | ***Hymd*** | ***Mat*** | *Raib* | *Rait* | *Asp* | ***Aspa*** | *Gon* | *Neo* | *Phy* | *Pro* | *Pte* | ***Tri*** |
| <u>MAR</u><br>(LI) | *Rr* | | | 54. 2% [34-73]<br>(3.4 ± 4.3) | | | | | | | | | 4.2% [0-20]<br>(0.1 ± 0.6) | | |
| <u>DIK</u><br>(LI) | *Rr* | 37.5% [20-58]<br>(6.7 ± 13.3) | | 25% [11-46]<br>(1 ± 1.8) | | | | | | 8.3% [1-27] | | | 4.2% [0-20]<br>(0.1 ± 0.2) | | |
| <u>DIT</u><br>(LI) | *Rr* | | | 59. 3% [40-76]<br>(3.8 ± 4.2) | 3.7% [0-18]<br>(0.2 ± 1) | | | 3.7% [0-18]<br>(1.1 ± 5.6) | | | | 3.7% [0-18]<br>(0.1 ± 0.3) | | | |
| <u>TOB</u><br>(LI) | *Rr* | | | 65% [42-83]<br>(4.2 ± 4.6) | | | | 5% [0-24]<br>(0.1 ± 0.4) | | | | 5% [0-25]<br>(0.1 ± 0.2) | | | |
| BOU<br>(IF) | *Rr* | | 3.4% [2-17]<br>(0.1 ± 0.2) | 3.4% [2-17]<br>(0.1 ± 1) | | | | | | | 3.4% [2-17]<br>(0.1 ± 0.2) | | | | |
| KED<br>(IF) | *Rr* | | | 41% [22-62]<br>(2.4 ± 4.3) | | | 4.5% [0-22]<br>(0.1 ± 0.21) | | | | 4.5% [0-22]<br>(0.2 ± 0.8) | 4.5% [0-22]<br>(0.1 ± 0.2) | | | |
| BAN<br>(IF) | *Mae* | | | | | | | | | | 11.1% [1-44]<br>(0.1 ± 0.3) | 11% [1-44]<br>(0.1 ± 0.3) | | | |
| BOU<br>(IF) | *Mae* | | | | | | | | | | 14.3% [1-55]<br>(0.3 ± 0.4) | | | | |
| BRA<br>(IF) | *Mae* | | | | | | | | | | | | | | 33.3% [2-86]<br>(1.7 ± 0.78) |
| KED<br>(IF) | *Man* | | 8.3% [1-27]<br>(0.1 ± 0.5) | | | 8.3% [1-27]<br>(0.2 ± 0.7) | 16.7% [6-37]<br>(1.4 ± 4.9) | | 12.5% [3-31]<br>(0.7 ± 2.9) | | 4.2% [0-20]<br>(0.1 ± 0.4) | | | | 45.8% [27-66]<br>(3.3 ± 5.1) |
| <u>BRA</u><br>(NI) | *Man* | | | | 54.5% [34-74]<br>(1.6 ± 1.9) | | | | | | | | | | 18.2% [6-39]<br>(1.1 ± 2.6) |
| <u>MAK</u><br>(NI) | *Man* | | | 3.8% [0-19]<br>(0.1 ± 0.2) | 23.1% [11-42]<br>(0.6 ± 1.4) | | | | 50% [30-63]<br>(12 ± 29.5) | | 3.8% [0-19]<br>(0.1 ± 0.2) | 3.8% [0-19]<br>(0.1 ± 0.4) | | 3.8% [0-19]<br>(0.1 ± 0.4) | 19.2% [8-38]<br>(0.4 ± 0.9) |
| <u>SEG</u><br>(NI) | *Man* | | | | 14.3% [4-35]<br>(0.5 ± 1.3) | | | | 38.1% [20-60]<br>(15.9 ± 65.6) | | 29.6% [13-51]<br>(1.0 ± 2.3) | | | | 52.4% [31-72]<br>(1.8 ± 3.9) |

Multivariate analyses revealed distinct helminth assemblages between native and invasive rodents (Figs. 2b, 3b), even on invasion fronts (Figs. 2a, 3a).

**Figure 2.**
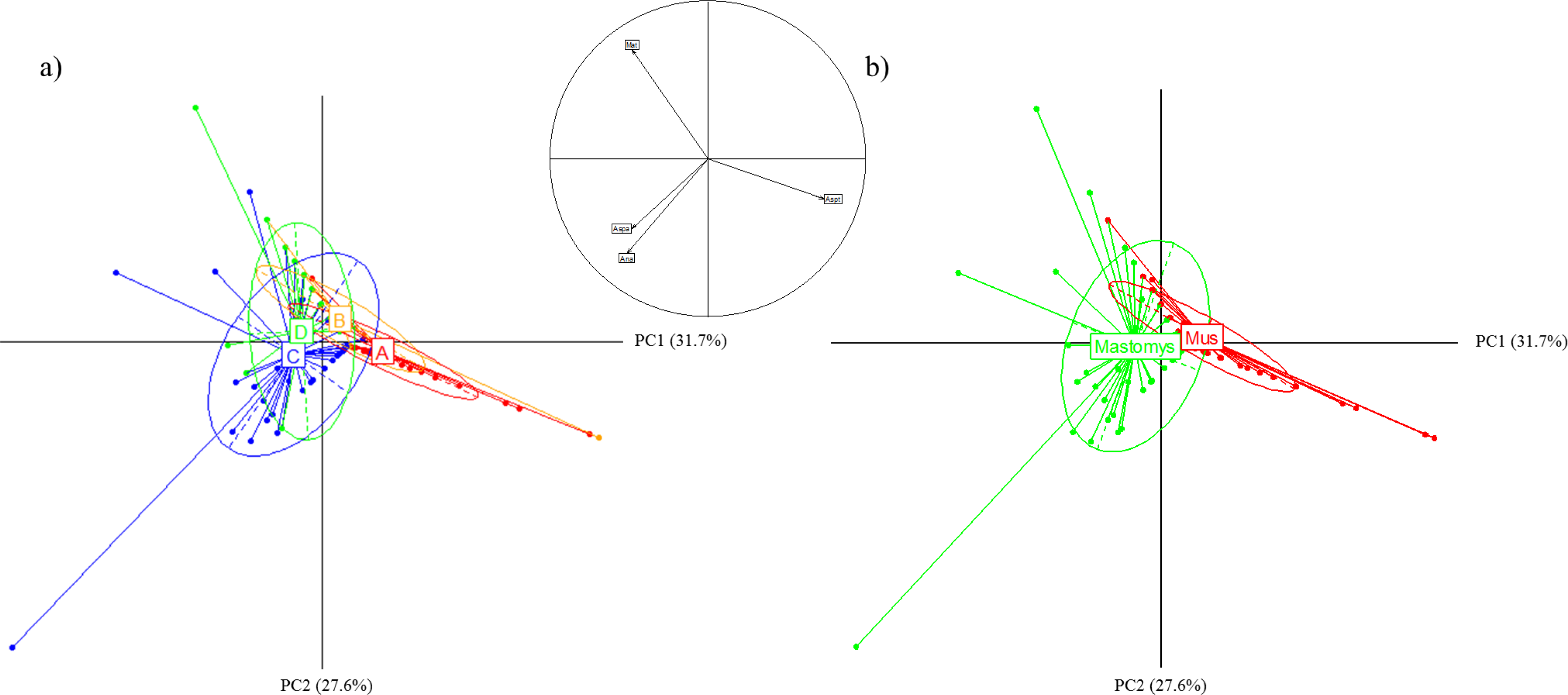
Principal component analysis showing GIH assemblages structure based on (a) the invasion status of the locality and (b) the host species on the mouse invasion route. Between-within analysis showed significant structuration for both factors (Monte-Carlo test, p < 0.05). The variables (GIH taxa having overall prevalence > 10%) are projected on the correlation circle between the two graphs (the codes used refer to those from table 2). Legend: A: *Mus (Mus musculus domesticus)* of long-established sites (red); B: Mus on invasion front (orange); C: *Mastomys* (*Mastomys erythroleucus*) on invasion front (blue); D: *Mastomys* in non-invaded sites (green).

**Figure 3.**
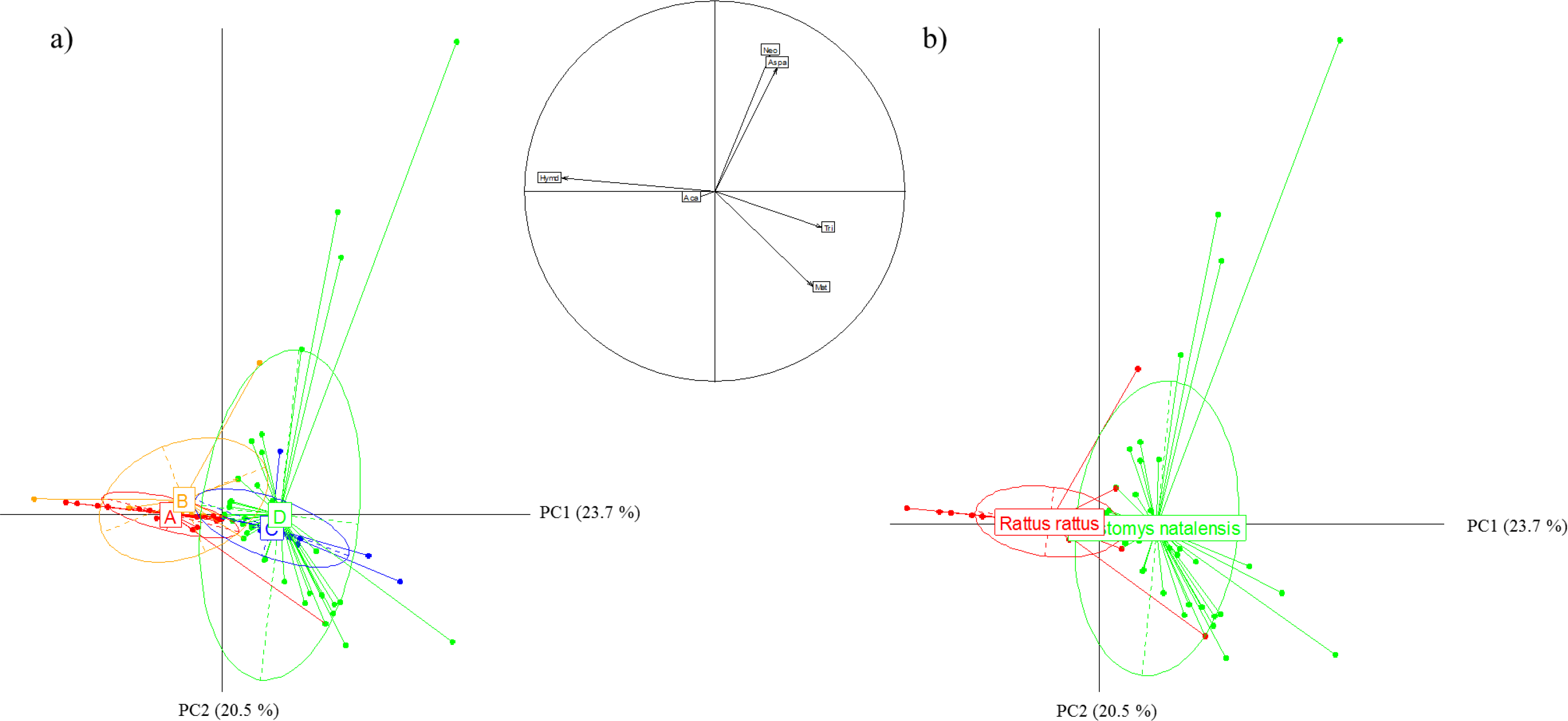
Principal component analysis showing GIH assemblages structure based on (a) the invasion status of the locality and (b) the host species on the rat invasion route. Between-within analysis showed significant structuration for both factors (Monte-Carlo test, p < 0.05). The variables (GIH taxa of which overall prevalence > 10%) are projected on the correlation circle between the two graphs (the codes used refer to those from table 3). This analysis considered only *Ma. natalensis* as native species because of too low GIH prevalence in *M. erythroleucus*. Legend: A: *Rattus rattus* of long-established rats (red); B: *R. rattus* on invasion front (orange); C: *Mastomys erythroleucus* on invasion front (blue); D: *Ma. erythroleucus* in noninvaded sites (green).

### 3.3 Testing factors affecting helminth assemblages

***Test of the enemy release hypothesis*** - For *M. m. domesticus*, GLMs revealed a significant effect of invasion status on helminth overall prevalence (F_1,166_ = 23.19, *p* < 0.0001), species richness (F_1,166_ = 25.22, *p* < 0.0001), and *Asp. tetraptera* prevalence (F1466 = 48.71, *p* < 0.0001), with higher values in host populations from long-established sites than in those from invasion front (Fig. 4; Table 4a).

**Figure 4.**
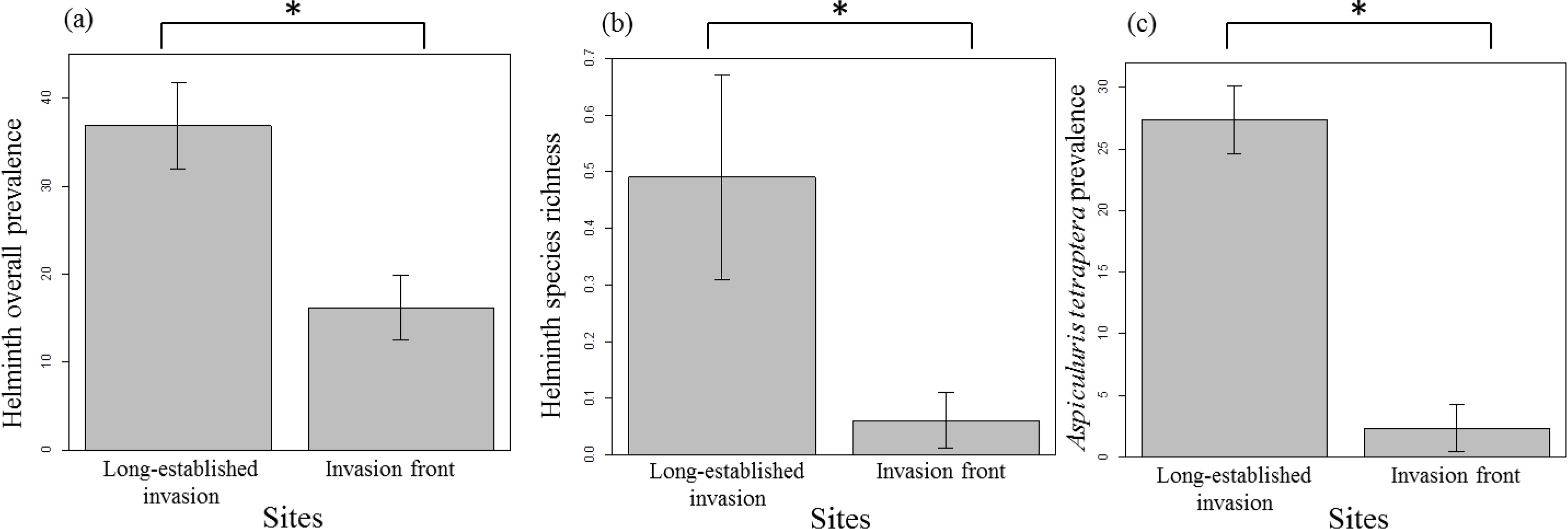
Difference in (a) helminth overall prevalence (presence/absence combining all taxa), (b) helminth species richness (number of taxa in one individual host), and (c) *Aspiculuris tetraptera* prevalence between house mouse populations from sites of long-established invasion (n = 95 individuals) and those from invasion front (n = 173).

**Table 4:**
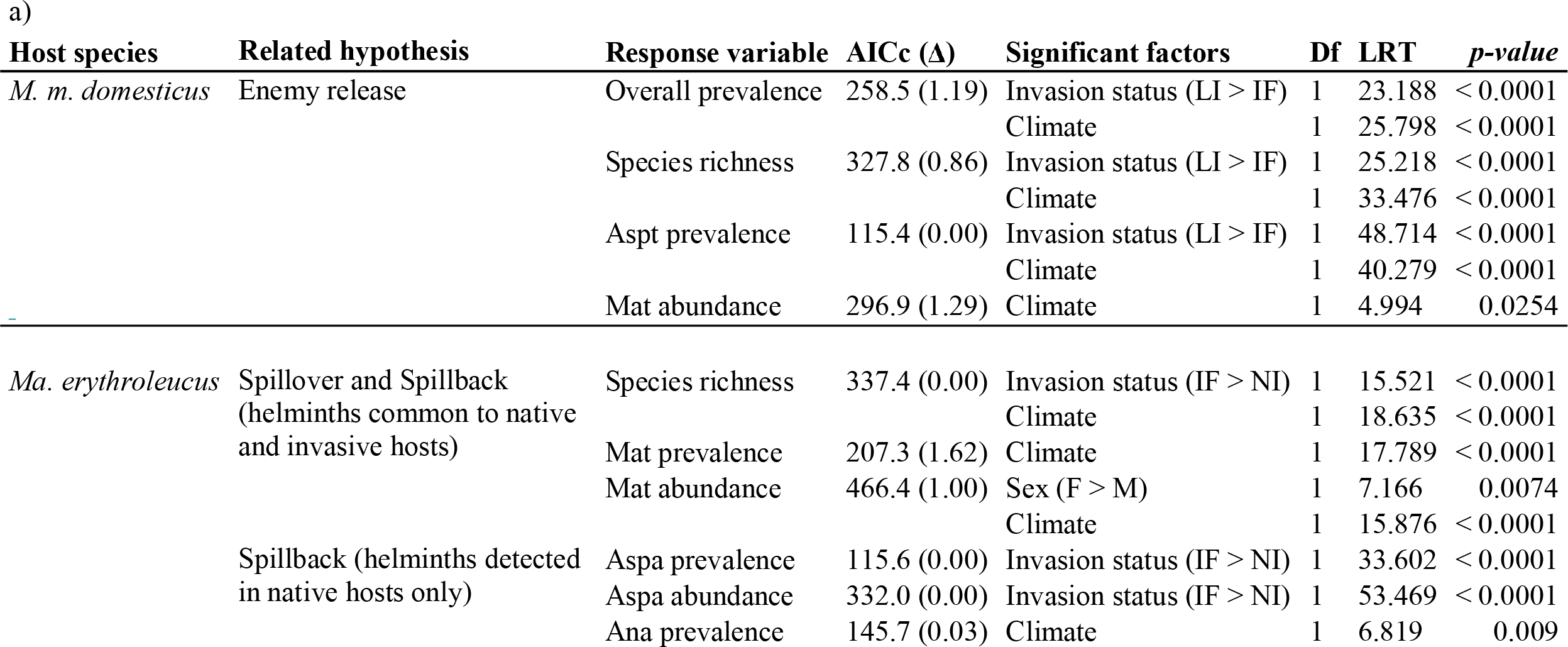
Most parsimonious Generalized Linear Models (GLMs) for the a) mouse and b) rat invasion routes. AlCc: Akaike’s information criterion corrected for finite sample size. Δ: difference between the model chosen and the model with the lowest AlCc. LRT: Likelihood-ratio test. LI: localities of long- established invasion; IF: invasion front; NI: non-invaded localities. F: Females; M: Males. Aspa: *Aspiculuris africana*; Aspt: *Aspiculuris tetraptera*; Mat: *Mathevotaenia* sp.; Tri: *Trichuris mastomysi*.

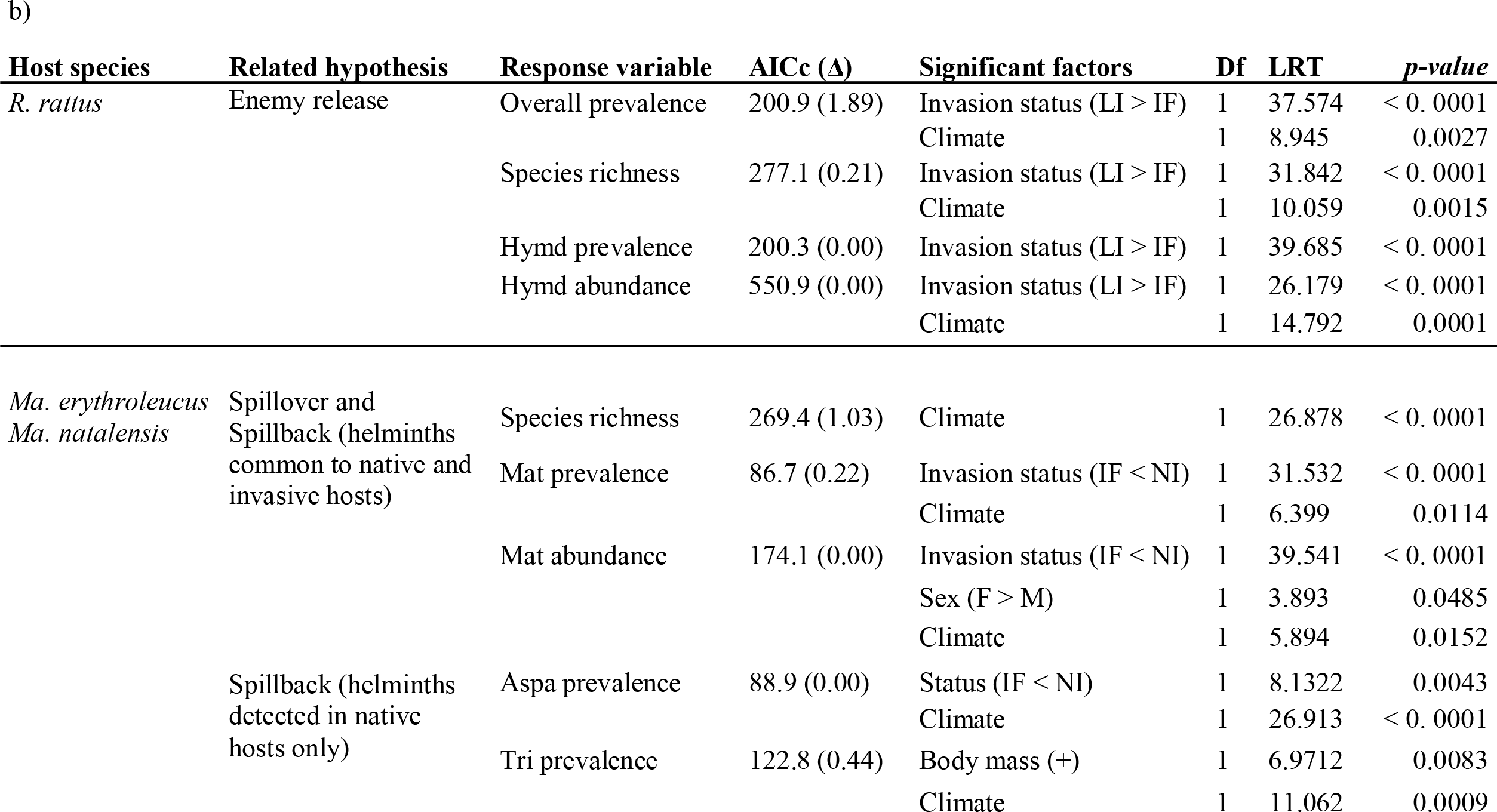

For *R. rattus,* GLMs revealed that helminth overall prevalence (F_1,191_ = 37.57, *p* < 0. 0001) and species richness (F_1,191_ = 31.84, *p* < 0. 0001) as well as *H. diminuta* prevalence (F_1,191_ = 39.69, *p* < 0. 0001) and abundance (F_1,191_ = 26.18, *p* < 0. 0001) were lower in sites of long-established invasion than in those at the invasion front (Fig. 5; table 4b).

**Figure 5.**
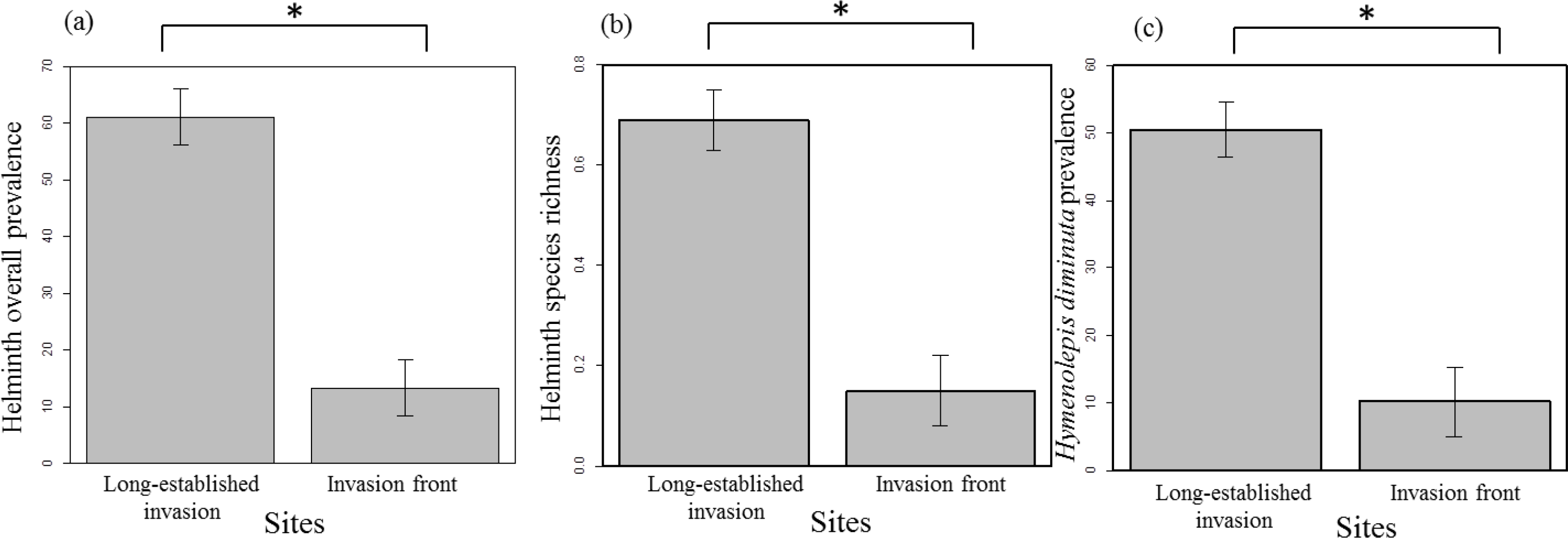
Difference in (a) helminth overall prevalence (presence/absence combining all taxa), (b) helminth species richness (number of taxa in one individual host), and (c) *Hymenolepis diminuta* prevalence between black rat populations from sites of long-established invasion (n = 95) and those from invasion front (n = 98).

These results were expected under the enemy-release hypothesis.

***Test of the spillover and spillback hypotheses*** - Native and invasive rodents shared some helminth parasites (e.g., the cestode *M. symmetrica* along the ‘mouse’ invasion route; the nematodes *Neoheligmonella granjoni, Physaloptera* sp., and the cestodes *H. diminuta, H. sp., M. symmetrica* and *R. trapezoides* along the ‘rat’ invasion route), which could be involved in spillover or spillback processes.

Along the ‘mouse’ invasion route, helminth species richness was higher for *Ma. erythroleucus* in sites of the invasion front than in not invaded sites (*F*_1,166_ = 25.22, *p* < 0.0001). Contrary to the expectations of spillover and spillback hypotheses, there was however no relationship between invasion status and prevalence or abundance of M *symmetrica* in *Ma erythroleucus.* Along the ‘rat’ invasion route, no relationship was found between helminth species richness and invasion status in *Mastomys* rodents. The cestode *M. symmetrica* was the only helminth common to native rodents and *R. rattus* with prevalence levels higher than 10% in native rodents. Contrary to spillover and spillback expectations, we found that *M. symmetrica* prevalence (*F*_1,120_ = 31.53, *p* < 0. 0001) and abundance (*F*_1,120_ = 39.54, *p* < 0. 0001) were lower at the invasion front than in non-invaded localities (Table 4b) in *Mastomys* spp.

***Effect of invasion status on helminths found specifically in native rodents*** - Following the results obtained for spillover and spillback hypotheses, we also made supplementary models on helminth taxa that were found only in native rodents (*Asp. africana* and *T. mastomysi*). These taxa may have been undetected in invasive rodents because of their low prevalence, which may lead to miss their involvement in spillback processes. In accordance with this expectation, the prevalence (*F*_1,167_ = 33.60, *p* < 0.0001) and abundance (*F*_1,167_ = 53.47, *p* < 0.0001) of *Asp. africana* were found to be higher in *Ma. erythroleucus* of the invasion front compared to those of non-invaded sites along the 'mouse’ invasion route (Table 4a). On the contrary, the prevalence of *Asp. africana* was found to be lower in native rodents of the invasion front compared to those of non-invaded sites along the ‘rat’ invasion route (*F*_1,120_ = 8.13, *p* = 0.0043; Table 4b).

***Effect of biological and climatic factors on helminth assemblages*** - Climate was included in several of the most parsimonious models (Table 4a, b). In exotic rodents, climate was found to explain helminth overall prevalence (*M. m. domesticus: F*_1,165_ = 25.80, *p* < 0.0001; *R. rattus: F*_1_,_190_ = 8.95, *p* = 0.0027), species richness (*M. m domesticus: F*_1,165_ = 33.48, *p* < 0.0001; R. *rattus: F*_1,190_ = 10.06, *p* = 0.0015), prevalence of *Asp. tetraptera* in M m. *domesticus* (F[,_26_5 = 40.28, *p* < 0.0001) and abundance of *M. symmetrica* in *M. m domesticus* (*F*_1,265_ = 4.99, *p* = 0.0254) or of *H. diminuta* in *R. rattus* (*F*_1,190_ = 14.79, *p* = 0.0001).

In native rodents, climate was found to explain the prevalence (*F*_1,166_ = 17.79, *p* < 0.0001) and abundance (*F*_1,165_ = 15.88, *p* < 0.0001) of *M. symmetrica,* and the prevalence of *Anatrichosoma* sp. (F1465 = 6.82, *p* < 0.009) along the ‘mouse’ invasion route, and species richness (*F*_1,120_ = 26.88, *p* < 0. 0001), M *symmetrica* prevalence (*F*_1,120_ = 6.40, *p* = 0.0114) and abundance (*F*_1,120_ = 39.54, *p* < 0. 0001), and *Anatrichosoma* sp. prevalence (*F*_1,165_ = 6.82, *p* < 0.009) along the ‘rat’ invasion route. Finally, females were more highly infected by *M. symmetrica* than males in native rodents along both the 'mouse’ (*F*_1_,_167_ = 7.17, *p* = 0.0074) and ‘rat’ (*F*_1,120_ = 3.89, *p* = 0. 0485) invasion routes.

## 4. Discussion

### Host specificity of helminth assemblages

In this study, we found nineteen helminth taxa in four rodent species. This high diversity is consistent with previous studies (e.g., Brouat *et al.* 2007; Elshazly *et al.* 2008). We found a small number of helminth taxa shared between native and exotic hosts. Along the ‘mouse’ invasion route, the only parasite shared was *M. symmetrica.* To date, this cestode was reported from invasive *R. rattus* and *M. m domesticus* in America, Europe and Asia (Beveridge 2008), but its recent identification in the South-African endemic rodent *Micaelamys namaquensis* (V. Haukisalmi, personal observation) makes the identification of its origin status less obvious. The large predominance of this species in *Mastomys* spp. along the ‘rat’ invasion route would support its native origin. Along the ‘rat’ invasion route, native and exotic hosts shared four helminth taxa, of which two, firstly described on African rodents (*N. granjoni* : Durette-Desset et al. 2008; *Raillietina trapezoides:* ref), are presumably native. The two other (*Hymenolepis* sp.; *Physaloptera* sp.), for which biological materials did not allow need to be identified at the species level.

The concomitant use of morphological and molecular tools gives confidence in the fact that several helminth taxa are host specific, and allows assessing their status as exotic or native parasites. Hence, *A. tetraptera* was found in several populations of *M. m. domesticus,* but not in native rodents. This nematode is known as a typical parasite of *M. m. domesticus* worldwide (Behnke *et al.* 2015), and would thus have been introduced by its host in Senegal. In the same way, *H. diminuta* was regularly found in *R. rattus,* but very occasionally (one infected individual) in native rodents. This cestode is retrieved in many regions of the world where *R. rattus* was introduced and was probably brought with its host (Elshazly *et al.* 2008). At last, *Asp. africana* and *T. mastomysi* were found exclusively in *Mastomys*. These nematodes were first described in African rodents (Verster, 1960; Quentin, 1966) and should thus be local parasites. Some helminth taxa found in this study exclusively in exotic rodents are otherwise reported in African natives, which makes the issue of their origin (native or exotic) less obvious. This is the case for *S. obvelata,* which is reported from various house mouse populations elsewhere (e.g., Milazzo *et al.* 2010), but also in local non-commensal rodents *Arvicanthis niloticus* and *Mastomys huberti* in Senegal (Diouf, 1994). Also, *P. senegalensis* found exclusively in *R. rattus* were first described in African rodents and may thus be a native parasite acquired by the exotic host.

### Weak pattern related to spillover and spillback hypotheses

Under the hypotheses of SO or SB, we expected to detect an increase of helminth species richness in native rodents from the invasion fronts compared to those from invaded sites, and an increase of specific prevalence or abundance of helminths shared between native and invasive hosts. These expectations were partially supported for native rodents along the ‘mouse’ invasion route, for helminth species richness and abundance of M *symmetrica.* Providing that this cestode is native, it may have been acquired by invasive rodents at the time of their establishment in coastal sites before spread. In this case, the increased infection level in *Ma. erythroleucus* at the invasion front would thus be a pattern compatible with the SB hypothesis.

On the rat invasion route, some helminth taxa were recorded in a rodent species exclusively at the invasion front, such as *Hymenolepis* sp. and *R. trapezoides* in *R. rattus* and *Ma. natalensis, R. baeri* in *M. natalensis* or *N. granjoni* in *R. rattus* (Table 3). Nevertheless, these taxa occurred in only a few sites with low infection levels and were not therefore considered in GLMs. These data indicated that host acquisition may have occurred at the invasion front as expected under SO/SB hypotheses. However, competence of the new host may be too low for the parasite acquisition has an impact on infection levels in native hosts. Moreover, lower infection levels of M *symmetrica* observed in *Mastomys* populations from the invasion front compared to non-invaded sites are not consistent with SO or SB hypotheses, and would rather suggest a dilution of this parasite (Johnson *et al.* 2008) when both host species occurred.

One can imagine that the detection of SO or SB patterns along mouse and rat invasion routes would be prevented by the involvement of highly virulent parasites causing the rapid death of the new host acquiring them. However, this argument is not likely for helminths that are generally not lethal for their hosts (Bordes & Morand 2011). The absence of typical SO or SB patterns may rather lie in the fact that many GIH taxa could be specialists or, at least, exhibit high host preferences, this latter potentially explaining the contrasted specific prevalence recorded for several helminths found in different rodent species (Tables 3, 4).

### Evidence for enemy loss

Many previous studies of parasites in invasive species may have over-estimated enemy release as they did not restrict comparison to the invasive population and the source populations from which it was founded (Colautti et al. 2004; Slothouber Galbreath et al. 2010). In the current study, to test for enemy release, whilst taking into account the source of invasion, the comparison of helminth diversity, prevalence and abundance was conducted only between current expanding populations and their well-defined source populations matched by molecular, historical and longitudinal data. Our data provided strong evidence that both *M. m. domesticus* and *R. rattus* exhibited lower rates of parasitism in sites sampled at the invasion front compared to long-established sites in Senegal, consistently with the enemy release hypothesis. This pattern was detected when considering the whole helminth community (overall prevalence and species richness) as well as specific taxa (*Asp. tetraptera* prevalence in *M. m. domesticus* and H *diminuta* prevalence and abundance in *R. rattus).* Enemy loss was already shown in expanding populations of *R. rattus* in other invasion contexts (Morand *et al.* 2015). To our knowledge, our work is the first providing such evidence of parasite loss in *M. m. domesticus.* Subsequently, our results raise obvious questions about the precise causes and outcomes of this parasite loss.

The decrease of parasitism levels in invading rodent populations may be explained either because the limited number of host individuals involved in spread does not carry the complete range of parasites found in source sites (founder effect), or because parasites that spread with their host are unable to establish and persist in the new environment of the invasion front (MacLeod *et al.* 2010). Distinguishing between these two types of processes is often difficult mainly because data on host and parasites close before and after the invasion spread are often lacking (Lymbery *et al.* 2014).

Particular features have been identified to render some parasites more prone to be lost during invasion (MacLeod *et al.* 2010). For instance, rare, patchily distributed or strongly virulent parasites have less opportunity to follow their host during its spread. Also, parasites with complex life cycles may fail to establish in a novel area because of sub-optimal environmental factors such as the absence of one of their required intermediate hosts. Helminths have usually low pathogenic effects due to co-evolution of immunoregulatory processes with their hosts (Dobson & Foufopoulos 2001), suggesting that strong virulence is not a key trait to explain their loss by invasive rodents. In this study, the two helminths that have been lost during mouse and rat expansions, i.e., *Asp. tetraptera* and *H diminuta* respectively, were highly prevalent in sites of long-established invasion. However, the absence of *Asp. tetraptera* in some of these sites (THL, NDB) may explain why this nematode is less prevalent at the invasion front of *M. m. domesticus.* Also, this directly transmitted parasite whose transmission depend on host density might have been brought by first introduced *M. m. domesticus* at the invasion front but then lost because of small host population size (Lippens *et al.* in revision). On the contrary, *H. diminuta* was homogeneously distributed in long-established invasion sites of *R. rattus.* The life cycle of this cestode requiring intermediate hosts might explain its loss during rat invasion. However, its intermediate hosts may be various arthropod species including beetles (Andreassen *et al.* 2004) that are *a priori* widely distributed in Senegal (Sembène *et al.* 2008).

Contrasted environmental conditions between sites of long-established invasion and invasion front may also explain spatial variations in helminth prevalence either directly (as helminths of terrestrial mammals spend at least one part of their life-cycle in the external environment outside their host) or indirectly through their impact on host demography or life-history traits (Krasnov *et al.* 1998). Consistently, climate has been systematically found to be significant in most of the models explaining variations in infection levels or community structure of helminths on both invasion routes.

Nevertheless, parasite loss does not necessarily mean parasite release (Prior *et al.* 2015). The extent to which parasite loss actually translates into a competitive advantage remains difficult to demonstrate because it involves subtle and complex impacts (Marcogliese & Pietrock 2011). For instance, a decrease in parasite species richness -as expected when PR occurs- may theoretically lead to lower inter-specific competition within parasite infracommunities, and thus increased abundance of the remaining parasites (Roche *et al.* 2010) or higher occurrence of over-regulated parasites within host populations. Assuming higher impacts of remaining parasites, this loss phenomenon should not have any positive effect on the host population. The loss of common parasites, such as *A. tetraptera* in *M. m. domesticus* or H *diminuta* in *R. rattus,* is more likely to result in an effective “release” for their host (Colautti *et al.* 2004). It has been advocated that enemy release may lead to positive outcomes in host either through regulatory (release of a parasite regulating host demographic parameters such as survivorship and fecundity) and/or through compensatory (reallocation of resources from defense to population growth over ecological time, or counter-selection of genotypes with costly defenses during invasion over evolutionary time) pathways (Colautti *et al.* 2004). Understanding how parasite loss may translate into effective release requires therefore a better understanding of the effects of specific enemies on their host (Colautti *et al.* 2004). Up to now, the advantage conferred by a parasite loss has more often been assumed than concretely addressed in animal models (Prior *et al.* 2015). Some studies provided field evidence that GIH may impact host population dynamics (Hudson 1998; Albon *et al.* 2002; Newey *et al.* 2005; Vandegrift & Hudson 2009; Rosà *et al.* 2011). Previous laboratory studies suggested relatively low effects of *Asp. tetraptera* on laboratory house mouse probably due to the selection of an immunological resistance (Derothe *et al.* 1997), but also negative effects on fitness in the case of heavy worm burdens (reviewed in Taffs, 1976). For *H. diminuta,* experimental infestations suggested expensive immunological responses accompanied with pathophysiological changes in hosts (Kosik-Bogacka, Baranowska-Bosiacka & Salamatin 2010). However, these studies focused on laboratory models that could differ from field-caught rodents in many ways. Further experimental infestations on M *m. domesticus* and *R. rattus* individuals sampled in natural populations in Senegal should be actually the best way to ascertain the helminth release-related benefices at the host population level, even if we are aware that the specific effects of a parasite in natural populations - as co-infections with other parasite taxa may occur in host infracommunities - could differ from those in common garden conditions (Telfer *et al.* 2010).

## Acknowledgments

We thank Laurent Granjon and Jean-Marc Duplantier for their precious help and advice during this work. Jean-Claude Berges and Natacha Volto are thanked for their help with climatic variables. We acknowledge the ANR ENEMI project (ANR-11-JSV7-0006) for funding and the French Embassy in Senegal for Ph.D. scholarships. We are particularly indebted to all the Senegalese people who allowed us to trap rodents in their homes.

---

**Supplementary Fig. S1.** Molecular phylogenetic tree of nematode sequences used as reference sequence database. The tree construction (Tamura-Nei model, Gamma distribution) was based on the mitochondrial Cytochrome Oxidase subunit 1 (CO1) following maximum-likelihood analysis with 100 bootstrap replicates implemented via the software MEGA6 (Molecular Evolutionary Genetics Analysis version 6.0). All worm samples were retrieved from rodent hosts collected in Senegal, Mali or Burkina Faso. Samples are identified by the host species and the code used to refer it in our collection. A number was added to “sp” when more than one undetermined species were expected for a particular nematode genus. Outgroup is sequences of the ac acanthocephalan Moniliformis moniliformis collected from R. rattus in Senegal. Scores at nodes represent bootstrap support for that node. Scale bar is proportional to the genetic distance in substitutions per site. Legend: A. *niloticus*: *Arvicanthis niloticus; Ma. erythroleucus*: *Mastomys erythroleucus; Ma. huberti: Mastomys huberti; Ma. natalensis: Mastomys natalensis; M. m. domesticus: Mus musculus domesticus; P. daltoni: Praomys daltoni; P. rostratus: Praomys rostratus; R rattus: Rattus rattus*.

**Supplementary Fig. S2**. Principal component analysis (PCA) of climatic data for categories of sites sampled along the house mouse invasion route (b) based on the uncorrelated climatic variables (temperatures in °C, rainfall in mm, recorded between 1997 and 2012) remaining after a first PCA (a). Between-within analysis showed significant classes (Monte-Carlo test, p < 0.05). Legend: max rain: maximum monthly rainfall during rainy season (mean per year); mn MnTM: lowest monthly minimum temperature (mean per year); sites of long-established invasion (red); sites of invasion front (blue); non-invaded sites (green). Temperature data were recorded from local weather stations closest to sampled sites and available on http://www.ncdc.noaa.gov/cdo-web/datasets; rainfall data were recorded from satellite products available on http://richardis.univparis1.fr/precip/rainser1.html with GPCP-1DD as date source.

**Supplementary Fig. S3**. Principal component analysis (PCA) of climatic data for categories of sites sampled along the black rat invasion route (b) based on the uncorrelated climatic variables (temperatures in °C, rainfall in mm, recorded between 1997 and 2012) remaining after a first PCA (a). Between-within analysis showed significant classes for both factors (Monte-Carlo test, p < 0.05). Legend: max MMxT: highest daily maximum temperature (mean per year); min MMnT: lowest daily minimum temperature (mean per year); min Rain: minimum monthly rainfall during rainy season (mean per year); sites of long-established invasion (red); sites of invasion front (blue); non-invaded sites (green). Temperature data were recorded from local weather stations closest to sampled sites and available on http://www.ncdc.noaa.gov/cdo-web/datasets; rainfall data were recorded from satellite products available on http://richardis.univparis1.fr/precip/rainser1.html with GPCP-1DD as date source.

